# Diagnostic Accuracy of FluoroCycler® XT MTBDR Assay for Detection of Rifampicin and Isoniazid Resistant *Mycobacteria tuberculosis* in Clinical Isolates from Kenya

**DOI:** 10.1101/2023.05.17.541083

**Authors:** Zakayo Maingi Mwangi, Samson Ireri, Haron Opwaka, Leon Otieno, Joan Simam, Frank Gekara Onyambu, Nellie Mukiri

**Author notes:** Corresponding author Mr. Zakayo Maingi Mwangi, Meru University of Science and Technology, P. O. Box: 972‑60200, Meru, Kenya. E‑mail. Corresponding author Nellie Mukiri, National Tuberculosis Reference Laboratory-Kenya P.O Box 20750 -00202, Nairobi, Kenya.

## Abstract

**Background:** Drug-resistant TB (DR-TB) poses a major global challenge to public health and therapeutics. It is an emerging global concern associated with increased morbidity and mortality mostly seen in the low- and middle-income countries. Lack of adequate diagnostic equipment for detection and monitoring of DR-TB leads to delayed diagnosis and subsequent inappropriate treatment. TB drug resistance testing has relied on phenotypic presentations in drug sensitivity testing (DST). The cost of setting up a TB phenotypic testing facility is prohibitive for most healthcare facilities due to its intensive investment in infrastructure, equipment, laboratory consumables, and personnel.

Molecular techniques are highly sensitive and offer timely and accurate results for TB drug resistance testing, thereby positively influencing patient management plan. The commonly used assay for detection of rifampicin (RIF) and isoniazid (INH) resistance in *Mycobacterium tuberculosis* (M.tb) is GenoType MTBDR*plus*. Although the GenoType MTBDR*plus* is more inexpensive and accurate than DST, when compared to other molecular techniques, it requires more specialized expertise, more hands-on time, substantial laboratory infrastructure and result interpretation is subjective to user. The FluoroCycler® MTBDR is a real-time polymerase chain reaction assay that detects M.tb and at the same time identifies mutations in *rpoB, katG and inhA* genes that are associated with RIF and INH resistance. It can detect up to 45 mutations in these genes in a single tube, producing results within 2.5 hours and this ability is only comparable to sequencing.

**Methods:** The study was carried out at the National Tuberculosis Reference Laboratory (NTRL) in Kenya in the period between January to October 2022. A total of 243 M.tb clinical isolates were included in the study. These isolates comprised of 50 isolates with mutations in *rpoB*, 51 isolates with *katG* mutations, 51 isolates with mutations in *inhA*. and 91 M.tb isolates lacking mutations in these genes based on Genotype MTBDR*plu*s results. DNA from the isolates was extracted using the FluoroLyse extraction kit. Real-time PCR targeting the *rpoB, InhA*, and *katG* genes was performed using the FluoroType MTBDR amplification mix. Isolates with discordant results between Genotype MTBDR*plu*s and FluoroCycler® MTBDR assays underwent targeted sequencing for the respective genes, then sequences were analyzed for mutations using Geneious version 11.0 software.

**Results:** The sensitivity of the Fluorocycler XT MTBDR assay for detection of mutations that confer drug resistance was 86% (95% CI 73.0,94.0) for *rpoB*, 96% (95% CI 87, 100) for *katG* and 92% (95% CI 81, 98) for *inhA*. The assay’s specificity was 97% (95% CI 93, 99) for *rpoB*, 98% (95% CI 96, 100) for *katG* and 97% (95% CI 93, 99) for *inhA*. Discrepancy between Genotype MTBDR*plus* and FluoroType MTBDR results were observed in 28 (11.5%) isolates with *rpoB, katG* and *inhA* genes having 26% (13/50), 10% (5/50), and 20% (10/50) isolates with discrepant results respectively. Sequencing results that were in agreement with FluoroType MTBDR results were 77% (10/13) for *rpoB*, 80% (4/5) for *katG*, and 70% (7/10) for *inhA* compared to 23% (3/13), 20% (1/5), and 30% (3/10) for Genotype MTBDR*plus* assay

**Conclusion:** The diagnostic accuracy of FluoroType MTBDR for the detection of mutations conferring resistance to RIF and INH was high compared with that of Genotype MTBDR*plus*, and demonstrates its suitability as a replacement assay for Genotype MTBDR*plus*.

## Introduction

Approximately 10 million people have TB with a total of 1.5 million deaths being recorded worldwide (WHO, 2022). Although the global incidence of TB is declining, Africa is disproportionately affected and contributes 2.6 million cases of the recorded global infections (Ismail et al., 2018). Together with other African countries, Kenya is among the high burden countries listed in all the three-list defined by WHO for TB, TB/HIV, and multidrug-resistant TB (MDR TB) where each category accounts for over 80% of the cases annually (Zignol et al., 2018). Drug resistant TB (DR-TB) is an emerging global concern and is associated with increased morbidity and mortality mostly observed in low- and middle-income countries (LMIC). Lack of adequate diagnostic equipment for detection and monitoring of DR-TB in LMIC causes delayed diagnosis and consequent treatment with ineffective anti-TB drugs.

Rifampicin (RIF) and isoniazid (INH) form the core of first-line anti-TB drugs and resistance to these two drugs account for more than 90% of MDR-TB (Meaza et al., 2021). Phenotypic characterization of *Mycobacterium tuberculosis* (M.tb) through culture and drug sensitivity testing (DST) are used to assess the drugs’ efficacy. Although DST is the gold standard for testing antibiotic efficacy, in Kenya, testing is done only at the National TB Reference Laboratory (NTRL) and select private healthcare facilities in the country, making the service less accessible to majority of patients. Mycobacterial culture and sensitivity require substantial investments in infrastructure, laboratory supplies, and manpower hence making it an expensive and prohibitive endeavor for majority of healthcare centers in LMIC. Further, DST has a long turnaround time (TAT) of eight weeks for culture and an additional two weeks for DST leading to delayed intervention and prolonged suffering for drug resistant TB cases (Cegielski et al., 2021). These diagnostic challenges have been addressed by the introduction of molecular assays which act by amplifying portions of the drug target genes, allowing for simultaneous detection of M.tb and identification of mutations linked to the development of drug resistance for RIF and INH in a shorter TAT. Molecular assays are suitable for most healthcare settings since they are more affordable, have a shorter TAT, and are highly sensitive compared to DST (Danfodiyo, 2020).

The molecular techniques that are approved so far by WHO for DR-TB testing include MTBDR*plus* (*Hain* Lifescience, Nehren, Germany) and Xpert MTB/RIF (Cepheid, Sunnyvale, CA). The Genotype MTBDR*plu*s (*Hain* Lifescience, Nehren, Germany) is a line probe assay (LPA) which employs the reverse hybridization method where amplicons are hybridized on a nitrocellulose membrane strip and result interpretation is done based on the presence or absence of bands at various points on the strip enabling for the simultaneous detection of M.tb and mutations associated with drug resistance for RIF and INH(Cegielski et al., 2021). The Xpert MTB/RIF (Cepheid, Sunnyvale, CA) is a real-time polymerase chain reaction (PCR) which amplifies and detects M.tb and mutations in the drug target gene for RIF only (World Health Organization, 2014).

The detection of M.tb and the diagnosis of RIF resistance at healthcare facilities and TB referral laboratories have been considerably enhanced by Xpert MTB/RIF and LPA assays which have subsequently improved TB management efforts and played a major role in reducing patient suffering(Enos et al., 2018). However, the LPA and Xpert MTB/RIF are biased in their detection of dominant mutations within the drug target genes and fail to capture silent mutations which also could contribute to drug resistance [5][6]. For instance, the RNA polymerase β subunit (*rpoB*) is the target gene for RIF, and 95% of RIF-resistant M.tb strains present a mutation within a 81bp dominant resistance determining region (RDR) which occurs between codons 507 to 533 of the gene. The other 5% of mutations are silent and occur outside the RDR but still contribute to RIF resistance as seen with DST. Also, 75% of INH drug resistance occurs following mutations in the RDR of *katG* and *inhA* genes which encodes the mycobacterial catalase-peroxidase and the InhA protein respectively, that are involved in mycolic acid synthesis (Karagoz et al., 2020). The non-RDR portions not considered in the above-mentioned analysis may contribute to the non-effectiveness of TB drugs (Tilahun et al., 2020; Zignol et al., 2018). Thus, the non-RDR for the drug target genes should also be evaluated together with the RDR portions for a comprehensive overview of the antimicrobial action of RIF and INH.

The FluoroCycler XT (*Hain* Lifesciences Nehren, Germany) is a molecular analyzer that utilizes the liquid array technology and relies on the Bruker-Hain Liquid Array technology that identifies *Mycobacteria tuberculosis* complex (MTBC) species, and detects mutations in *rpoB, inhA* and *katG* genes (De Vos et al., 2018; Dippenaar et al., 2021). It captures both dominant and silent mutations in *rpoB, katG*, and *inhA* genes thereby yielding a detailed report on the nature of drug resistance if any for each drug. The assay can detect up to 45 mutations in these genes within one tube and produces results in 2.5 hours and this ability is only comparable to sequencing. The sensitivity of this assay for detecting MTBC is enhanced by its high limit of detection identifying as low as 14 colony forming units/milliliter (CFU/ML) and 8 CFU/ML for culture and sputum samples respectively (Vos et al., 2018).

We therefore, evaluated the accuracy of FluoroCycler® XT MTBDR assay in comparison to MTBDR*plus* in the detection of mutations within *rpoB, katG* and *inhA* genes.

## Materials and methods

The study was carried out at the National Tuberculosis Reference Laboratory (NTRL) in Kenya in the period between January to October 2022. NTRL receives sputum samples from patients for TB and DR-TB testing using culture and MTBDR*plu*s respectively. A total of 243 M.tb clinical isolates, with both culture and MTBDR*plu*s results, were included in the study. These isolates comprised of 50 isolates with mutations in *rpoB*, 51 isolates with *katG mutations*, 51 isolates with mutations in *inhA*. and 91 M.tb isolates lacking mutations in these genes based on Genotype MTBDR*plu*s results were included.

### DNA extraction

DNA was extracted from cultured isolates using a FluoroLyse kit (Hain Lifescience GmbH) according to the manufacturer’s instructions. Briefly, a 500μl aliquot of the harvested colonies was centrifuged for 15 min at 10,000 x *g* to pellet the bacilli. The resulting pellet was resuspended in 100 μl lysis buffer (F-LYS) and mixed with 2 μl of internal control then vortexed for 5 seconds. The mixture was then incubated for 5 min at 95°C. Subsequently, 100 μl of the neutralization buffer (F-NB) was added, and the mixture was vortexed for 5 seconds and then centrifuged for 5 min at 14,000 x *g*. The supernatant was then transferred to a new tube and stored at -20°C until further use. A tube containing no clinical material was included during the extraction protocol as a negative control.

PCR mixes were freshly prepared by combining 6 μl amplification mix A (AM-A) and 14 μl amplification mix B (AM-B). Thereafter, 20 μl of the FluoroLyse-extracted DNA was added to the PCR mix. Positive control provided in the Fluorotype kit and negative control prepared during the DNA extraction process were also included at 20 μl for each. The PCR mixes were immediately loaded into the FluoroCycler96 instrument to avoid photobleaching.

### Analysis

For determination of FluoroCycler XT accurac*y* in detection of RIF and INH resistance, the FluoroType results were compared to the Genotype MTBDR*plus* hybridization patterns of the probes corresponding to the *rpoB* and *katG* genes and the *inhA* promoter region. Sensitivity, specificity, negative predictive value (NPV) and positive predictive value (PPV) were calculated for ***rpoB, inhA*, and *katG*** separately. Sensitivity was calculated as the percent proportion of isolates determined as resistant by FluoroCycler XT compared to Genotype MTBDR*plus*. Specificity was calculated as the percent proportion of isolates determined as susceptible by FluoroCycler XT compared to MTBDR*plus*, PPV was calculated as the percent proportion of isolates determined as resistant by FluoroCycler XT among isolates identified as resistant by Genotype MTBDR*plus*. NPV was calculated as the percent proportion of susceptible isolates determined by FluoroCycler XT among isolates identified as susceptible by Genotype MTBDR*plus*. Cohen’s kappa coefficient was used to determine the degree of agreement between FluoroCycler XT and MTBDR*plus* (Fleiss, 1981). Kappa values indicate as follows: 0 shows no agreement, 0.10-0.20 slight agreement, 0.21-0.40-fair agreement, 0.41-0.60 moderate agreement, 0.61-0.80 substantial agreement, 0.81-0.90 near perfect agreement and 1 indicates perfect agreement. To compare the proportion of resistant isolates identified FluoroCycler XT and Genotype MTBDR*plus*, we used McNemar test on paired proportions where discordant pairs between the two assays was greater than 10. The statistical analysis was performed using R Statisitcal software (v3.6.3; R Core Team 2020)). Results with P value < 0.05 were considered as statistically significant.

For isolates with discrepant results between MTBDR*plus* and FluoroCycler XT, Sanger sequencing was applied as a reference method to ascertain the true status of the drug target genes.

### Targeted sequencing of *rpoB, katG*, and *inhA for samples with discordant results*

Samples with discordant results underwent conventional PCR for the respective genes using the Horse-Power™ Taq DNA Polymerase MasterMix (Canvax, Córdoba, Spain) in a final reaction volume for each gene of 13μl comprising 6.25μl of 2X Horse-Power™ Taq DNA Polymerase MasterMix, 2.5μl DNA template, 0.25μl of each of both forward and reverse primers (Table 1) at a final concentration of 10 pmoles, and 3.75μl of nuclease-free water to make up the reaction volume. The PCR assays were carried out with a Veriti Thermal Cycler (Applied Biosystems, Foster City, CA, USA).

**Table 1:**
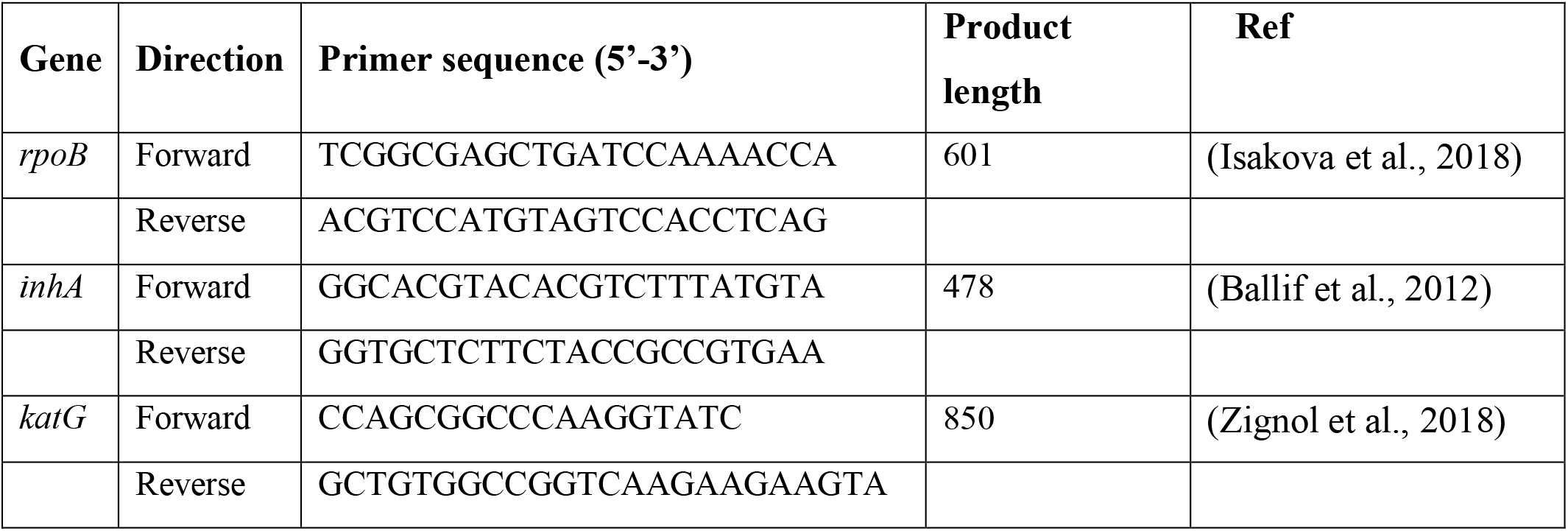
Primer sequence for *rpoB, inhA*, and *katG*

Thermal cycling conditions for *rpoB* were as follows: 1 cycle of 95°C for 5 minutes, 35 cycles of 95°C for 1 minute, 62°C for 1 minute, 72°C for 1 minute, and a final extension for 10 minutes at 72°C. PCR for *kat G* was conducted as follows 95°C for 5 minutes, 35 cycles of 95°C for 1 minute, 64°C for 1 minute, 72°C for 1 minute, and a final extension for 7 minutes at 72°C. PCR for the *inhA* was conducted as follows: 95°C for 5 minutes, 35 cycles of 95°C for 1 minute, 60°C for 1 minute, 72°C for 1 minute, and a final extension for 7 minutes at 72°C. Amplified products were confirmed on a 1% Agarose gel stained with 4.6μl SYBR safe DNA stain (Invitrogen, Carlsbad, California, USA), and results were visualized with an UltraViolet gel viewer (Terra Universal, <u>S. Raymond Ave, Fullerton, CA</u>,USA).

The PCR products were enzymatically purified using ExoSAP IT (Applied Biosystems, Foster City, California, USA). Purification conditions were done at 37ºC for 15 minutes followed by a second incubation at 80ºC for 15 minutes and a final cooling step at 4ºC for 5 minutes.

The purified amplicons were sequenced in the forward and reverse directions by Sanger sequencing using Big Dye™ Terminator Version 3.1 Cycle Sequencing Kit (Applied Biosystems, Foster City, California, USA) and the forward and reverse primers. The sequencing reaction for each gene was a 10μl reaction comprising 1.25 μl of Big Dye Terminator, 3 μl of 5X Sequencing Buffer, 1 μl of 1 pmol of the sequencing primer, and 1.5 μl of the PCR product. The reaction volume was made up by adding 3.25 μl of nuclease-free water. The reaction proceeded through 96°C for 1 minute then 25 cycles of 96°C for 10 seconds, 50°C for 5 seconds, and 60°C for 4 minutes.

Purification of cycle-sequencing products was done using the BigDye X Terminator™ purification kit following the manufacturer’s instructions (Applied Biosystems, Foster City, California, USA) and purified products were loaded onto the ABI 3730 genetic analyzer (Applied Biosystems, Foster City, California, USA) for capillary electrophoresis. The obtained sequences were aligned to their wild-type reference sequences for each gene using Geneious version 11.0 (Biomatters Ltd, Auckland, New Zealand). Mutations in the drug resistance genes were identified visually.

## Results

The sensitivity of the Fluorocycler XT MTBDR assay for detection of mutations that confer drug resistance was; *rpoB* 86% (95% CI 73.0, 94.0), *katG* 96% (95% CI 87, 100) and *inhA* 92% (95% CI 81, 98) while the specificity was: *rpoB* 97% (95% CI 93, 99), *katG* 98% (95% CI 96, 100) and *inhA* 97% (95% CI 93, 99). The positive and negative predictive values ranged between 88%-99%. There was a near perfect agreement between the two assays witih Kappa values greater than 0.84. The proportion of resistant isolates identified by the two assays do not statistically differ by McNemar Test (p=0.97) (Table 2).

**Table 2:** **Diagnostic accuracy of the FluoroCycler® XT MTBDR V2.0 Assay for detection of mutations in *rpoB, katG* and, *inhA* genes in comparison to Genotype** MTBDR*plus* assay.

| Gene | FluoroCycler® XT |  | Genotype MTBDR<sub>plus</sub> |  | Sensitivity (95%CI) | Specificity (95% CI) | PPV (95% CI) | NPV (95% CI) | Kappa Statistic | McNemar (P value) |
| --- | --- | --- | --- | --- | --- | --- | --- | --- | --- | --- |
|  | R | S | R | S |  |  |  |  |  |  |
| *rpoB* | 49 | 194 | 50 | 193 | 86% (73.0,94.0) | 97% (93, 99) | 96 (93, 99) | 96 (93, 99) | 0.835 | 0.97(1) |
| *katG* | 52 | 191 | 51 | 192 | 96% (87, 100) | 98% (96, 100) | 94 (84, 99) | 99 (96, 100) | 0.878 | - |
| *inhA* | 53 | 190 | 51 | 192 | 92% (81, 98) | 97% (93, 99) | 89 (77, 96) | 88 (75, 95) | 0.938 | 0.97(0.7518) |

\*R-Resistant S-Sensitive

Using **Fluorotype MTBDR**, majority of mutations within *rpoB* were observed at position 531(48%, n=22) with base substitution occurring at S531F (48%, n=10), S531L (31.8%, n=7), S531Q (18.2%, n=4), S531W (4.5%, n=1). Additional mutation in *rpoB* included deletions at position 517 in seven isolates (15%) and at position 517-519 in one isolate (2%), substitutions at D516F/Y (13%, n=6), H526L/N (8%, n=4), L533D (4%, n=2), and *combined mutation at position Q513H and D516Y (4%, n=2), N518L (2%, n=1), L511P (2%, n=1) S522L (2%, n=1) (Table 3).

**Table 3:**
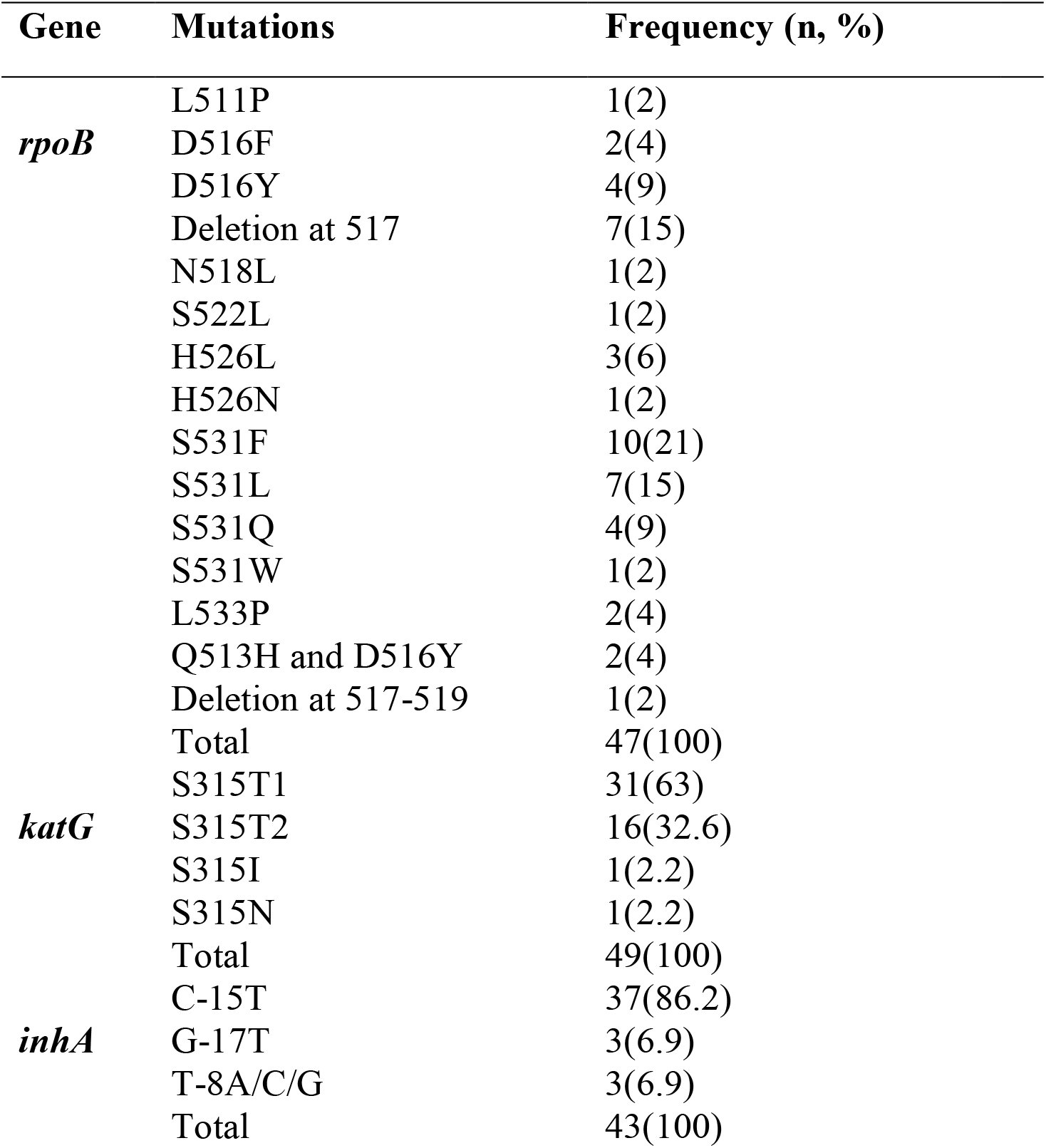
Mutation patterns within the *rpoB, katG* and *inhA* genes isolates identified by Fluorotype MTBDR

The *katG* gene had four forms of mutations at position 315 including S315T1 (n=31, 63%) and S315T2 (n=16, 32.6%), S315I (n=1, 2.2%) and S315N ((n=1, 2.2%) (Table 3). Three loci within *inhA* had base substitutions with 86.2% (37/43) having a C-15T mutation, 6.9% (3/43) at G-17T and 6.9% (3/43) having T-8A/C/G mutations (Table 3).

Discrepancy between Genotype MTBDR*plus* and FluoroType MTBDR results were observed in 28 (11.5%) isolates. FluoroType MTBDR detected *rpoB, katG* and *inhA* mutations in 6, 3 and 6 isolates respectively that were identified as susceptible by Genotype MTBDR*plus* assay. Point mutations not detected by Genotype MTBDR*plus* but captured by FluoroType MTBDR include V517E, S522L, G523G, and H526L/N/Q in *rpoB*, S315I/R/N in *katG*, and T-17C, T-8G/C, T-15C, and T-8C in *inhA* Sanger sequencing was used to address the discrepancy. Sequencing results were in agreement with FluoroType MTBDR in 21/28 (75%) isolates whereas Genotype MTBDR*plus* assay was in agreement in 7/28 (25%) isolates (Table 4). In addition, FluoroType MTBDR showed a higher agreement with sequencing identification of mutation in the three genes. We observed agreement in 10/13 (77%), 4/5 (80%), and 7/10) (70%) for *rpoB, katG and inhA* respectively isolates compared to 3/13 (23%), 1/5 (20%), and 3/10) (30%) Genotype MTBDR*plus* assay

**Table 4:** Resolution of discrepant results for mutations in *rpoB, katG* and *inhA* genes for Genotype MTBDR*plus* and FluoroType MTBDR using Sanger sequencing

| Gene | Genotype MTBDR<sub>plus</sub> | Fluorotype MTBDR | Sanger sequencing |
| --- | --- | --- | --- |
| *rpoB* | WT | V517E | WT |
|  | WT | S522L | WT |
|  | WT | H526L | WT |
|  | MUT-2A | WT | WT |
|  | MUT-2A | WT | WT |
|  | MUT-2B | WT | WT |
|  | MUT-2A | WT | WT |
|  | MUT-2B | WT | WT |
|  | MUT-2A | WT | WT |
|  | MUT-2A | WT | WT |
|  | WT | H526L | H526L |
|  | WT | H526N | H526N |
|  | WT | S531Q | S531Q |
| *katG* | WT | S315I | WT |
|  | MUT 2 | WT | WT |
|  | MUT 2 | WT | WT |
|  | WT | S315T | S315R |
|  | WT | S315N | S315R* |
| *inhA* | *WT | C-15T | *WT |
|  | MUT1 | WT | WT |
|  | MUT1 | WT | WT |
|  | MUT2 | WT | A-16G |
|  | MUT2 | WT | A-16G |
|  | *WT | G-17T | *G-17T |
|  | *WT | T-8G | *T-8G |
|  | *WT | T-8C | *T-8C |
|  | *WT | T-8A | *T-8A |
|  | *WT | C-15T | *T-15C |

## Discussion

The accuracy of the FluoroType MTBDR assay for detection of mutations-conferring RIF resistance in *rpoB* was high with a sensitivity of 86% (95% CI 73.0, 94.0) and specificity of 98% (95% CI 96, 100) respectively when using Genotype MTBDR*plu*s assay. These results are similar to those of a study done using smear positive RIF resistant samples that showed a sensitivity and specificity of 99.3% (95.8–100) and 100 (97.2–100) respectively (Vos et al., 2018). The FluoroType MTBDR assay was able to correctly classify *rpoB* mutations according to their types. For instance, majority of mutations within *rpoB* were observed at position 531(48%, n=22) with base substitution occurring as S531F (48%, n=10), S531L (31.8%, n=7), S531Q (18.2%, n=4), S531W (4.5%, n=1). It also identified deletion at positions 517 (15%, n=7) and at position 517-519 (2%, n=1,), substitution at D516F/Y (13%, n=6), H526L/N (8%, n=4), L533D (4%, n=2), and combined mutation at position Q513H&D516Y (4%, n=2), N518L (2%, n=1), L511P (2%, n=1) S522L (2%, n=1), no base insertion mutation types were seen for our RIF resistant isolates. Identification of mutations types occurring in particular loci within *rpoB* is important since it would guide therapeutic strategies in precision medicine of gene mutation, after biological and functional mechanisms of the subsequent polymerase enzyme is clearly validated (Wang, 2016). The FluoroType MTBDR assay was also able to identify low-level resistance mutations in *rpoB* at position 511, 516, 522 and 533. These mutations exhibit elevated RIF minimum inhibitory concentrations (MICs) compared to fully susceptible strains by DST, but remain phenotypically susceptible and have been associated with poor patient outcomes (Shea et al., 2021). There were no silent mutations detected thus demonstrating the rarity of mutations outside the RDR.

In our investigation, the Fluorotype MTBDR assay’s ability to detect resistance for INH was done by examining the *katG* and *inhA* genes. The sensitivity and specificity of the assay in detecting mutations in *katG* gene was 96% (95% CI 87, 100) and 97% (95% CI 93, 99) respectively with most mutations occurring at position 315 (S315T/I/R,N) (63%, 31/49) which confers a high-level INH resistance with MICs ranging from 2 to >10 mg/L and can go up to 25.6 mg/L with DST (Hsu et al., 2020). The accuracy of Fluorotype MTBDR was comparable to a previous study that presented a sensitivity of 94.3% (85.3-98.2) and specificity of 99.5% (96.8-100) for the same gene (Dippenaar et al., 2021).

The FluoroType MTBDR assay’s sensitivity to mutations in *inhA* was 92% (95% CI 81, 98) while its specificity was 97% (95% CI 93, 99). This result was comparable to a previous study that showed 96% sensitivity (De Vos et al., 2018). A mutation limited only to the *inhA* promoter area is usually associated with low-level resistance and a higher dose of INH (10-15mg/kg/day), instead of the usual dose in first-line regimens (4-6mg/kg/day) (Liu et al., 2018).

Isolates with concordant results in resistance for Genotype MTBDR*plus* and FluoroType MTBDR were 86% (43/50) in *rpoB*, while *katG* and *inhA* were 96% (49/51) and 92% (47/51) respectively. While no significant difference in the performance between the two assays for the detection of RIF and INH resistance was observed, a higher discrepancy was observed between Genotype MTBDR*plu*s and Sanger sequencing compared to FluoroType MTBDR. There was a discrepancy in 18 isolates in detection of resistance and susceptibility between Genotype MTBDR*plu*s and Sanger sequencing as opposed to 9 isolates using FluoroType MTBDR assay. This could be due to reader subjectivity while visually interpreting the hybridization patterns in Genotype MTBDR*plu*s. For instance, point mutations not detected by Genotype MTBDR*plus* but captured by FluoroType MTBDR and confirmed by targeted sequencing include V517E, S522L, G523G, and H526L/N/Q in *rpoB*, S315I/R/N in *katG*, and T-17C, T-8G/C, T-15C, and T-8C in *inhA*.

## Ethical approval

This study was approved by MUST Institutional Research Ethics Review Committee (*MIRERC*) (Ref: MU/1/39/28 Vol.2:67)) on March 28, 2022. Waiver for individual informed consent was granted as the study utilized remnant clinical samples and the research posed no greater than minimal risk to the study subjects.

## Conclusion

The capacity of Fluorocycler MTBDR to concurrently detect a wide variety of mutations in the *rpoB, katG*, and *inhA* promoter may be helpful in directing specialized treatment plans. This confirms its applicability as a substitute assay for Genotype MTBDR*plu*s, without the subjectivity of visually evaluating the hybridization patterns. The results of the FluoroType assay are also independently assessed and presented by the analyzer software, as opposed to the Genotype MTBDR*plu*s assay, which needs human or semiautomated grading of hybridization patterns. An advantage of the FluoroType MTBDR assay is its performance in a single closed tube, preventing the release of amplicons, eliminating the possibility of cross-contamination in the lab, and requiring less infrastructure in the lab compared to Genotype MTBDR*plu*s or DST.

## Limitation

Hetero-resistance and indeterminate results were not carefully examined in our study, and could have contributed to the inconsistent results between the Genotype MTBDR*plus* and Fluorocycler MTBDR.

However, the objective of this study was to analyze the diagnostic accuracy of the Fluorocycler MTBDR and we concluded that its diagnostic capability of detecting RIF and INH resistance was higher compared to Genotype MTBDR*plus*, and that Fluorocycler MTBDR should be implemented as a replacement for Genotype MTBDR*plus* at the NTRL.

