## Supplemental Table 1 for "Diagnostic Accuracy of FluoroCycler® XT MTBDR Assay for Detection of Rifampicin and Isoniazid Resistant *Mycobacteria tuberculosis* in Clinical Isolates from Kenya"

| S/no | Sample ID | Gend | age | County | Type of patient | MTBDRplus | | | Fluorocycler XT | | | Sequencing | | |
|---|---|---|---|---|---|---|---|---|---|---|---|---|---|---|
| | | | | | | INH | KATG | RPOB | INH | KATG | RPOB | INH | KATG | RPOB |
| 1 | Fluor-001 | M | 48 | MOMBASA | New | S | R | S | S | R | S | | | |
| 2 | Fluor-002 | M | 31 | MERU | New | S | R | R | S | R | R | | | |
| 3 | Fluor-003 | M | 50 | MERU | New | R | S | S | R | S | S | | | |
| 4 | Fluor-004 | M | 35 | KIAMBU | New | S | S | S | S | S | R | | | S |
| 5 | Fluor-005 | M | 38 | TIGANIA | Retreatment | R | S | S | R | S | S | | | |
| 6 | Fluor-006 | M | 37 | NYANDARI | New | S | S | S | S | S | S | | | |
| 7 | Fluor-007 | F | 29 | NAIROBI | New | R | S | S | R | S | S | | | |
| 8 | Fluor-008 | M | 33 | NYERI | relapse | S | R | R | S | R | R | | | |
| 9 | Fluor-009 | M | 38 | NAIROBI | New | R | S | S | R | S | S | | | |
| 10 | Fluor-010 | F | 21 | KAJIADO | Retreatment | S | S | S | S | S | S | | | |
| 11 | Fluor-011 | M | 20 | NAKURU | New | S | S | R | S | S | R | | | |
| 12 | Fluor-012 | M | 31 | LAIKIPIA | New | S | R | S | S | R | S | | | |
| 13 | Fluor-013 | M | 40 | MURANGA | Retreatment | R | S | S | S | S | S | R | | |
| 14 | Fluor-014 | M | 59 | THARAKA | New | S | S | S | S | S | S | | | |
| 15 | Fluor-015 | M | 40 | MERU | Retreatment | S | S | S | S | S | S | | | |
| 16 | Fluor-016 | M | 40 | MERU | New | S | S | S | S | S | S | | | |
| 17 | Fluor-017 | F | 53 | NAKURU | New | R | S | R | S | S | S | S | | S |
| 18 | Fluor-018 | M | 59 | MACHAKO | Retreatment | S | R | S | S | R | S | | | |
| 19 | Fluor-019 | F | 36 | MOMBASA | MDR-fu | S | S | R | S | S | R | | | |
| 20 | Fluor-020 | M | 41 | MERU | MDR-fu | S | S | S | S | S | S | | | |
| 21 | Fluor-021 | F | 14 | BUSIA | New | S | S | R | S | S | R | | | |
| 22 | Fluor-022 | F | 30 | KIAMBU | relapse | R | S | R | S | S | R | S | | |
| 23 | Fluor-023 | M | 68 | MAKUENI | New | R | S | R | R | S | R | | | |
| 24 | Fluor-024 | M | 40 | GARISSA | New | S | S | S | S | S | R | | | R |
| 25 | Fluor-025 | M | 88 | WEST POK | New | R | R | S | R | R | S | | | |
| 26 | Fluor-026 | M | 23 | UASIN GISI | relapse | S | R | S | S | R | S | | | |
| 27 | Fluor-027 | M | 46 | MANDERA | New | S | S | S | S | S | S | | | |
| 28 | Fluor-028 | F | 30 | KIAMBU | relapse | S | S | S | S | S | S | | | |
| 29 | Fluor-029 | F | 22 | KAJIADO | relapse | R | R | S | R | R | S | | | |
| 30 | Fluor-030 | M | 22 | TURKANA | Retreatment | S | S | S | R | S | S | S | | |
| 31 | Fluor-031 | F | 18 | WEST POK | Retreatment | S | R | S | S | R | S | | | |
| 32 | Fluor-032 | M | 50 | TURKANA | relapse | S | S | R | S | S | R | | | |
| 33 | Fluor-033 | M | 27 | KILIFI | relapse | S | S | R | S | S | R | | | |
| 34 | Fluor-034 | M | 26 | MERU | New | R | S | S | R | S | S | | | |
| 35 | Fluor-035 | M | 33 | MARSABIT | Retreatment | S | S | S | S | S | S | | | |
| 36 | Fluor-036 | M | 42 | KIAMBU | relapse | S | S | S | S | S | S | | | |
| 37 | Fluor-037 | M | 40 | MERU | relapse | S | S | S | S | S | S | | | |
| 38 | Fluor-038 | M | 48 | MERU | New | S | S | R | S | S | S | | | S |
| 39 | Fluor-039 | M | 48 | EMBU | New | S | R | S | S | R | S | | | |
| 40 | Fluor-040 | F | 46 | WEST POK | relapse | S | S | S | S | S | S | | | |
| 41 | Fluor-041 | F | 48 | NAIROBI | MDR-fu | S | S | S | S | S | S | | | |
| 42 | Fluor-042 | M | 35 | MACHAKO | Retreatment | S | S | R | S | S | S | | | S |
| 43 | Fluor-043 | M | 33 | MERU | New | R | S | R | R | S | R | | | |
| 44 | Fluor-044 | F | 24 | BUSIA | New | S | S | S | S | S | S | | | |
| 45 | Fluor-045 | F | 30 | MERU | New | S | S | S | S | S | S | | | |
| 46 | Fluor-046 | M | 40 | BARINGO | New | R | S | R | R | S | R | | | |
| 47 | Fluor-047 | M | 38 | NAIROBI | New | R | S | R | R | S | R | | | |
| 48 | Fluor-048 | F | 21 | MERU | New | S | S | S | S | S | S | | | |
| 49 | Fluor-049 | M | 41 | MERU | MDR-fu | R | S | S | S | S | S | S | | |
| 50 | Fluor-050 | M | 48 | KITUI | New | S | R | R | S | R | R | | | |
| 51 | Fluor-051 | F | 30 | MERU | MDR-fu | S | S | R | S | R | R | | R | |
| 52 | Fluor-052 | M | 40 | MURANGA | Retreatment | S | S | S | S | S | R | | | S |
| 53 | Fluor-053 | M | 20 | BUNGOMA | Retreatment | R | R | S | R | S | S | | S | |
| 54 | Fluor-054 | M | 38 | KAJIADO | New | S | S | S | S | S | S | | | |
| 55 | Fluor-055 | M | 33 | MARSABIT | Retreatment | S | R | R | S | R | R | | | |
| 56 | Fluor-056 | M | 38 | NAIROBI | NEW | R | S | S | R | S | S | | | |
| 57 | Fluor-057 | F | 29 | NAIROBI | New | S | S | R | S | S | R | | | |
| 58 | Fluor-058 | F | 32 | KISUMU | MDR-fu | S | S | S | S | S | S | | | |
| 59 | Fluor-059 | M | 45 | POKOT | New | S | S | S | S | S | S | | | |
| 60 | Fluor-060 | F | 45 | MERU | New | S | S | S | R | S | S | R | | |
| 61 | Fluor-061 | M | 35 | MACHAKO | Retreatment | R | S | S | R | S | S | | | |
| 62 | Fluor-062 | F | 46 | WEST POK | relapse | S | S | S | S | S | S | | | |
| 63 | Fluor-063 | F | 44 | KERICHO | New | S | S | R | S | S | R | | | |
| 64 | Fluor-064 | M | 38 | MERU | relapse | S | S | S | S | S | S | | | |
| 65 | Fluor-065 | M | 20 | MANDERA | New | S | S | S | S | S | S | | | |
| 66 | Fluor-066 | M | 35 | NAKURU | relapse | R | S | S | R | S | S | | | |
| 67 | Fluor-067 | M | 53 | NAKURU | relapse | R | S | S | R | S | S | | | |
| 68 | Fluor-068 | M | 19 | NAKURU | New | R | R | R | R | R | R | | | R |
| 69 | Fluor-069 | M | 47 | NANDI | relapse | S | S | S | S | S | S | | | |
| 70 | Fluor-070 | M | 46 | TRANS NZ( | New | S | S | R | S | S | S | | | S |
| 71 | Fluor-071 | M | 19 | KITUI | relapse | S | S | R | S | S | R | | | |
| 72 | Fluor-072 | M | 35 | NAKURU | relapse | S | R | R | S | R | R | | | |
| 73 | Fluor-073 | M | 31 | LAIKIPIA EA | New | R | R | R | R | R | R | | | |
| 74 | Fluor-074 | M | 26 | MERU | New | S | S | S | S | S | S | | | |
| 75 | Fluor-075 | M | 33 | MOMBASA | New | S | R | S | S | R | S | | | |
| 76 | Fluor-076 | M | 55 | BUSIA | New | S | R | S | S | R | R | | | S |
| 77 | Fluor-077 | M | 29 | MARSABIT | New | R | R | S | R | R | S | | | |
| 78 | Fluor-078 | M | 35 | EMBU | MDR-fu | S | S | R | S | S | R | | | |
| 79 | Fluor-079 | F | 16 | KILIFI | relapse | R | R | R | R | R | R | | | |
| 80 | Fluor-080 | M | 42 | KWALE | MDR-fu | S | S | S | S | S | S | | | |
| 81 | Fluor-081 | F | 39 | KIRINYAGA | New | R | R | S | R | R | R | | | R |
| 82 | Fluor-082 | M | 40 | MERU | relapse | S | S | R | S | S | R | | | |
| 83 | Fluor-083 | F | 30 | TURKANA | relapse | R | S | S | R | S | S | | | |
| 84 | Fluor-084 | M | 37 | EMBU | New | R | R | S | R | R | S | | | |
| 85 | Fluor-085 | F | 23 | EMBU | New | R | S | S | R | S | S | | | |
| 86 | Fluor-086 | M | 26 | KILIFI | relapse | R | S | R | R | S | R | | | |
| 87 | Fluor-087 | M | 40 | EMBU | New | R | S | S | R | S | S | | | |
| 88 | Fluor-088 | F | 26 | NAIROBI | relapse | S | S | S | S | S | S | | | |

| | | | | | | | | | | | | | | |
|---|---|---|---|---|---|---|---|---|---|---|---|---|---|---|
| 89 | Fluor-089 | F | 21 | KIAMBU | MDR-fu | S | S | S | S | S | S | | | |
| 90 | Fluor-090 | F | 22 | NAIROBI | MDR-fu | R | R | S | R | R | S | | | |
| 91 | Fluor-091 | F | 22 | NYANDARI | New | S | S | R | R | S | R | R | | |
| 92 | Fluor-092 | M | 36 | NAIROBI | New | S | S | R | S | R | R | | R | |
| 93 | Fluor-093 | F | 23 | EMBU | New | S | R | S | S | R | S | | | |
| 94 | Fluor-094 | F | 24 | MERU | New | S | R | S | S | R | S | | | |
| 95 | Fluor-095 | M | 38 | MERU | relapse | S | R | S | S | R | S | | | |
| 96 | Fluor-096 | M | 45 | POKOT | New | S | S | R | S | S | R | | | |
| 97 | Fluor-097 | M | 29 | MARSABIT | New | R | R | S | R | R | S | | | |
| 98 | Fluor-098 | M | 31 | NAKURU | New | R | S | R | R | S | R | | | |
| 99 | Fluor-099 | F | 21 | KAJIADO | Retreatment | S | S | R | S | R | R | | S | |
| 100 | Fluor-100 | M | 23 | TURKANA | relapse | S | S | S | S | S | S | | | |
| 101 | Fluor-101 | F | 28 | BARINGO | New | S | R | S | S | R | S | | | |
| 102 | Fluor-102 | F | 29 | POKOT | New | S | R | R | S | R | R | | | |
| 103 | Fluor-103 | F | 30 | POKOT | New | R | R | S | R | R | S | | | |
| 104 | Fluor-104 | M | 50 | TURKANA | relapse | S | S | S | S | S | S | | | |
| 105 | Fluor-105 | M | 24 | MAKUENI | relapse | S | S | S | S | S | S | | | |
| 106 | Fluor-106 | M | 38 | MERU | Retreatment | S | S | R | S | S | R | | | |
| 107 | Fluor-107 | M | 33 | MACHAKO | Retreatment | S | R | R | S | R | R | | | |
| 108 | Fluor-108 | M | 50 | GARISSA | New | S | R | S | S | R | S | | | |
| 109 | Fluor-109 | M | 26 | EMBU | Retreatment | S | R | S | S | R | R | | | R |
| 110 | Fluor-110 | F | 29 | KITUI | MDR-fu | R | S | S | R | S | S | | | |
| 111 | Fluor-111 | F | 27 | GARISSA | New | S | S | R | S | S | R | | | |
| 112 | Fluor-112 | M | 63 | KITUI | New | S | S | S | S | S | S | | | |
| 113 | Fluor-113 | M | 50 | GARISSA | New | S | R | R | S | R | R | | | |
| 114 | Fluor-114 | M | 56 | MERU | New | S | R | R | R | S | S | S | S | S |
| 115 | Fluor-115 | F | 21 | KAJIADO | Retreatment | R | S | S | R | S | S | | | |
| 116 | Fluor-116 | M | 21 | MERU | New | S | S | S | S | S | S | | | |
| 117 | Fluor-117 | F | 31 | TRANS NZ( | New | S | S | S | S | S | S | | | |
| 118 | Fluor-118 | M | 30 | MERU | relapse | S | S | S | S | S | S | | | |
| 119 | Fluor-119 | M | 48 | EMBU | New | S | S | S | S | S | S | | | |
| 120 | Fluor-120 | F | 21 | KAJIADO | Retreatment | R | S | R | R | S | R | | | |
| 121 | Fluor-121 | M | 26 | MERU | relapse | R | S | S | R | S | S | | | |
| 122 | Fluor-122 | M | 65 | GARISSA | New | S | R | S | S | R | S | | | |
| 123 | Fluor-123 | M | 45 | KAKEMEG/ | relapse | S | S | S | R | S | S | R | | |
| 124 | Fluor-124 | M | 22 | TRANS NZ( | Retreatment | R | R | R | R | R | S | | | S |
| 125 | Fluor-125 | M | 28 | MERU | New | R | R | S | R | R | S | | | |
| 126 | Fluor-126 | F | 21 | UASIN GIS | Retreatment | S | S | R | R | S | R | R | | |
| 127 | Fluor-127 | M | 52 | MERU | relapse | S | S | S | S | S | S | | | |
| 128 | Fluor-128 | M | 38 | GARISSA | relapse | S | R | R | S | R | R | | | |
| 129 | Fluor-129 | F | 31 | MOMBASA | New | R | R | R | R | R | S | | | S |
| 130 | Fluor-130 | F | 32 | MERU | New | R | R | S | R | R | S | | | |
| 131 | Fluor-131 | F | 32 | GARISSA | New | S | R | R | S | R | R | | | |
| 132 | Fluor-132 | M | 32 | GARISSA | New | S | R | S | S | R | S | | | |
| 133 | Fluor-133 | M | 31 | KILIFI | New | S | R | R | S | R | R | | | |
| 134 | Fluor-134 | M | 30 | KAJIADO | New | R | R | R | R | R | R | | | |
| 135 | Fluor-135 | M | 29 | NAIROBI | New | S | R | S | S | R | S | | | |
| 136 | Fluor-136 | M | 33 | KIAMBU | New | S | R | R | S | R | R | | | |
| 137 | Fluor-137 | M | 32 | KIAMBU | Relapse | S | R | S | S | R | S | | | |
| 138 | Fluor-138 | M | 31 | KILIFI | Relapse | R | R | S | R | R | S | | | |
| 139 | Fluor-139 | M | 29 | MERU | Relapse | S | R | R | S | R | R | | | |
| 140 | Fluor-140 | M | 27 | NYERI | Relapse | S | S | S | S | S | S | | | |
| 141 | Fluor-141 | M | 34 | KAKEMEG/ | Relapse | S | R | S | S | R | S | | | |
| 142 | Fluor-142 | M | 33 | NANDI | Relapse | R | S | S | R | S | S | | | |
| 143 | Fluor-143 | M | 38 | TURKANA | Relapse | S | S | S | S | S | S | | | |
| 144 | Fluor-144 | M | 37 | TURKANA | Relapse | S | S | S | S | S | S | | | |
| 145 | Fluor-145 | M | 41 | UASIN GIS | Retreatment | S | S | S | S | S | S | | | |
| 146 | Fluor-146 | M | 40 | BUNGOM/ | Retreatment | R | S | S | R | S | S | | | |
| 147 | Fluor-147 | F | 45 | TRANS NZ( | MDR-fu | S | S | S | S | S | S | | | |
| 148 | Fluor-148 | F | 53 | MACHAKO | MDR-fu | S | S | S | S | S | S | | | |
| 149 | Fluor-149 | M | 19 | MACHAKO | MDR-fu | S | S | S | S | S | S | | | |
| 150 | Fluor-150 | M | 20 | MACHAKO | MDR-fu | R | S | S | R | S | S | | | |
| 151 | Fluor-151 | F | 20 | MERU | New | S | S | S | S | S | S | | | |
| 152 | Fluor-152 | F | 21 | NAIROBI | New | S | S | S | S | S | S | | | |
| 153 | Fluor-153 | F | 26 | NYANDARI | New | S | S | S | S | S | S | | | |
| 154 | Fluor-154 | F | 26 | EMBU | New | R | S | S | R | S | S | | | |
| 155 | Fluor-155 | F | 28 | KAJIADO | New | S | S | S | S | S | S | | | |
| 156 | Fluor-156 | F | 29 | NAIROBI | New | S | S | S | S | S | S | | | |
| 157 | Fluor-157 | F | 29 | NAIROBI | New | S | S | S | S | S | S | | | |
| 158 | Fluor-158 | M | 31 | NAIROBI | New | S | S | S | S | S | S | | | |
| 159 | Fluor-159 | M | 31 | EMBU | New | R | S | S | R | S | S | | | |
| 160 | Fluor-160 | M | 31 | GARISSA | New | S | S | S | S | S | S | | | |
| 161 | Fluor-161 | M | 31 | MERU | New | S | S | S | S | S | S | | | |
| 162 | Fluor-162 | M | 33 | BARINGO | New | S | S | S | S | S | S | | | |
| 163 | Fluor-163 | M | 33 | POKOT | New | R | S | S | R | S | S | | | |
| 164 | Fluor-164 | M | 35 | BUSIA | New | S | S | S | S | S | S | | | |
| 165 | Fluor-165 | M | 36 | MERU | NEW | S | S | S | S | S | S | | | |
| 166 | Fluor-166 | M | 37 | THARAKA | New | S | S | S | S | S | S | | | |
| 167 | Fluor-167 | M | 37 | KITUI | New | S | S | S | S | S | S | | | |
| 168 | Fluor-168 | M | 38 | GARISSA | New | S | S | S | S | S | S | | | |
| 169 | Fluor-169 | M | 29 | MAKUENI | New | S | S | S | S | S | S | | | |
| 170 | Fluor-170 | M | 26 | WEST POK | New | S | S | S | S | S | S | | | |
| 171 | Fluor-171 | M | 30 | KILIFI | New | S | S | S | S | S | S | | | |
| 172 | Fluor-172 | M | 30 | KAJIADO | New | S | S | S | S | S | S | | | |
| 173 | Fluor-173 | M | 30 | MAKUENI | New | S | S | S | S | S | S | | | |
| 174 | Fluor-174 | M | 46 | KILIFI | New | S | S | S | S | S | S | | | |
| 175 | Fluor-175 | F | 46 | MERU | New | S | S | S | S | S | S | | | |
| 176 | Fluor-176 | F | 19 | KILIFI | New | S | S | S | S | S | S | | | |
| 177 | Fluor-177 | F | 23 | MERU | New | S | S | S | S | S | S | | | |
| 178 | Fluor-178 | F | 23 | NYERI | New | S | S | S | S | S | S | | | |

| | | | | | | | | | | | |
|---|---|---|---|---|---|---|---|---|---|---|---|
| 179 | Fluor-179 | M | 24 | NAKURU | New | S | S | S | S | S | S |
| 180 | Fluor-180 | M | 26 | NAKURU | relapse | S | S | S | S | S | S |
| 181 | Fluor-181 | M | 26 | GARISSA | relapse | S | S | S | S | S | S |
| 182 | Fluor-182 | M | 27 | MERU | relapse | S | S | S | S | S | S |
| 183 | Fluor-183 | M | 30 | MERU | relapse | S | S | S | S | S | S |
| 184 | Fluor-184 | M | 33 | MERU | relapse | S | S | S | S | S | S |
| 185 | Fluor-185 | M | 35 | NANDI | relapse | S | S | S | S | S | S |
| 186 | Fluor-186 | F | 35 | TURKANA | relapse | S | S | S | S | S | S |
| 187 | Fluor-187 | F | 38 | TURKANA | relapse | S | S | S | S | S | S |
| 188 | Fluor-188 | F | 38 | MERU | relapse | S | S | S | S | S | S |
| 189 | Fluor-189 | F | 38 | NAKURU | relapse | S | S | S | S | S | S |
| 190 | Fluor-190 | M | 40 | WEST POK | relapse | S | S | S | S | S | S |
| 191 | Fluor-191 | M | 40 | TRANS NZ | relapse | S | S | S | S | S | S |
| 192 | Fluor-192 | M | 42 | TURKANA | Retreatment | S | S | S | S | S | S |
| 193 | Fluor-193 | M | 45 | EMBU | Retreatment | S | S | S | S | S | S |
| 194 | Fluor-194 | M | 47 | MARSABIT | Retreatment | S | S | S | S | S | S |
| 195 | Fluor-195 | M | 30 | MACHAKO | Retreatment | R | S | S | R | S | S |
| 196 | Fluor-196 | M | 32 | MACHAKO | Retreatment | S | S | S | S | S | S |
| 197 | Fluor-197 | M | 36 | MERU | Retreatment | S | S | S | S | S | S |
| 198 | Fluor-198 | M | 48 | TIGANIA | Retreatment | S | S | S | S | S | S |
| 199 | Fluor-199 | F | 35 | MERU | New | S | S | S | S | S | S |
| 200 | Fluor-200 | F | 41 | MURANGA | New | S | S | S | S | S | S |
| 201 | Fluor-201 | F | 41 | MURANGA | New | S | S | S | S | S | S |
| 202 | Fluor-202 | F | 42 | BARINGO | New | S | S | S | S | S | S |
| 203 | Fluor-203 | F | 24 | POKOT | New | S | S | S | S | S | S |
| 204 | Fluor-204 | F | 21 | POKOT | New | S | S | S | S | S | S |
| 205 | Fluor-205 | F | 23 | MOMBASA | New | S | S | S | S | S | S |
| 206 | Fluor-206 | F | 24 | MERU | New | S | S | S | S | S | S |
| 207 | Fluor-207 | M | 27 | GARISSA | New | S | S | S | S | S | S |
| 208 | Fluor-208 | M | 29 | GARISSA | New | S | S | S | S | S | S |
| 209 | Fluor-209 | M | 29 | KILIFI | New | S | S | S | S | S | S |
| 210 | Fluor-210 | M | 30 | KAJIADO | New | S | S | S | S | S | S |
| 211 | Fluor-211 | M | 31 | NAIROBI | New | S | S | S | S | S | S |
| 212 | Fluor-212 | M | 39 | KIAMBU | New | S | S | S | S | S | S |
| 213 | Fluor-213 | M | 44 | NYERI | New | S | S | S | S | S | S |
| 214 | Fluor-214 | M | 45 | KAKEMEGA | relapse | S | S | S | S | S | S |
| 215 | Fluor-215 | M | 53 | NANDI | relapse | S | S | S | S | S | S |
| 216 | Fluor-216 | M | 19 | TURKANA | relapse | S | S | S | S | S | S |
| 217 | Fluor-217 | F | 20 | TURKANA | relapse | S | S | S | S | S | S |
| 218 | Fluor-218 | F | 20 | UASIN GIS | relapse | S | S | S | S | S | S |
| 219 | Fluor-219 | F | 21 | BUNGOMA | relapse | S | S | S | S | S | S |
| 220 | Fluor-220 | F | 26 | TRANS NZ | relapse | S | S | S | S | S | S |
| 221 | Fluor-221 | F | 26 | KITUI | Retreatment | S | S | S | S | S | S |
| 222 | Fluor-222 | M | 28 | MERU | Retreatment | S | S | S | S | S | S |
| 223 | Fluor-223 | M | 29 | KISUMU | Retreatment | S | S | S | S | S | S |
| 224 | Fluor-224 | M | 31 | MOMBASA | Retreatment | S | S | S | S | S | S |
| 225 | Fluor-225 | M | 28 | NAIROBI | Retreatment | S | S | S | S | S | S |
| 226 | Fluor-226 | F | 33 | EMBU | Retreatment | S | S | S | S | S | S |
| 227 | Fluor-227 | F | 35 | MERU | Retreatment | S | S | S | S | S | S |
| 228 | Fluor-228 | F | 36 | MERU | Retreatment | S | S | S | S | S | S |
| 229 | Fluor-229 | F | 37 | KWALE | Retreatment | S | S | S | S | S | S |
| 230 | Fluor-230 | F | 34 | BUSIA | Retreatment | S | S | S | S | S | S |
| 231 | Fluor-231 | F | 38 | MERU | Retreatment | S | S | S | S | S | S |
| 232 | Fluor-232 | F | 38 | MERU | Retreatment | S | S | S | S | S | S |
| 233 | Fluor-233 | F | 38 | TRANS NZ | Retreatment | S | S | S | S | S | S |
| 234 | Fluor-234 | F | 39 | KIRINYAGA | Retreatment | S | S | S | S | S | S |
| 235 | Fluor-235 | M | 40 | KERICHO | Retreatment | S | S | S | S | S | S |
| 236 | Fluor-236 | M | 40 | MERU | Retreatment | S | S | S | S | S | S |
| 237 | Fluor-237 | M | 40 | NAKURU | Retreatment | S | S | S | S | S | S |
| 238 | Fluor-238 | M | 46 | NAKURU | Retreatment | S | S | S | S | S | S |
| 239 | Fluor-239 | M | 45 | NAKURU | Retreatment | S | S | S | S | S | S |
| 240 | Fluor-240 | M | 45 | MANDERA | Retreatment | S | S | S | S | S | S |
| 241 | Fluor-241 | M | 44 | MERU | New | S | S | S | S | S | S |
| 242 | Fluor-242 | M | 46 | MERU | New | S | S | S | S | S | S |
| 243 | Fluor-243 | M | 21 | MERU | New | S | S | S | S | S | S |
